# A Multiscale Immuno-Oncology on-Chip System (MIOCS) establishes that collective T cell behaviors govern tumor regression

**DOI:** 10.1101/2021.03.23.435334

**Authors:** Gustave Ronteix, Shreyansh Jain, Christelle Angely, Marine Cazaux, Roxana Khazen, Philippe Bousso, Charles N. Baroud

**Affiliations:** Physical microfluidics and Bioengineering, Institut Pasteur, 25-28 Rue du Dr. Roux, Paris, France; LadHyX, CNRS, Ecole polytechnique, Institut polytechnique de Paris, 91120, Palaiseau, France; Dynamics of Immune Responses Unit, Equipe Labellisée Ligue Contre le Cancer, Institut Pasteur, INSERM U1223, 75015 Paris, France

## Abstract

T cell-based tumor immunotherapies such as CAR T cells or immune checkpoint inhibitors harness the cytotoxic potential of T cells to promote tumor regression. However, patient response to immunotherapy remains heterogeneous, highlighting the need to better understand the rules governing a successful T cell attack. Here, we develop a microfluidic-based method to track the outcome of T cell activity on many individual cancer spheroids simultaneously, with a high spatiotemporal resolution. By combining these parallel measurements of T cell behaviors and tumor fate with probabilistic modeling, we establish that the first recruited T cells initiate a positive feedback loop leading to an accelerated effector accumulation on the spheroid. We also provide evidence that cooperation between T cells on the spheroid during the killing phase facilitates tumor destruction. We propose that tumor destruction does not simply reflect the sum of individual T cell activities but relies instead on collective behaviors promoting both T cell accumulation and function. The possibility to track many replicates of immune-tumor interactions with such a level of detail should help delineate the mechanisms and efficacy of various immunotherapeutic strategies.

## I. INTRODUCTION

The capacity of cytotoxic T lymphocytes (CTLs) to eliminate tumor cells is the basis for the development of important tumor immunotherapies such as immune checkpoint inhibitors (e.g anti-CTLA-4, anti-PD1 or anti PD-L1 mAbs) and the development of cellular therapies such as CAR T cells [1, 2]. However, patient response to these therapies can be highly variable. While many parameters are known to influence patient response to immunotherapies, the number, phenotype and distribution of CTLs can have a strong predictive value in several types of cancer [3].

These observations underscore the need to better understand how a successful T cell attack proceeds and what are the critical parameters associated with CTL behavior and function that favor tumor regression. In this respect, several key questions remain unanswered. For example, how do CTLs encounter tumor cells and accumulate within the tumor microenvironment (TME)? What are the dynamics of CTL killing and what CTL density is needed for tumor eradication? Are individual CTLs acting autonomously in the TME or do they interact together? Understanding the basic principles that dictate whether a tumor mass is regressing or not is in fact essential to design, optimize and evaluate tumor immunotherapeutic strategies.

Multiple approaches are available to evaluate T cell cytotoxicity against tumors. *In-vitro* assays in cell suspension have been used to measure CTL killing capacity and, when performed at the single-cell level, provide information on the extent of functional heterogeneity within a T cell population [4]. These in vitro assays however lack the complexity of the 3D tumor microenvironment, which strongly impacts T cell behavior and function. At the other end of the spectrum, intravital imaging offers direct insights into the dynamics, signaling and killing behavior of single T cells within a developing tumor [5–8]. Limitations of these approaches however include the fact that they provide a view of the interactions in a limited spatial and temporal window. Indeed continuous observation periods are generally limited to a few hours, precluding a full understanding of T cell histories in the TME.

An interesting emerging platform comes from advanced *in-vitro* models that recapitulate some aspects of the TME while providing access to the system dynamics [9, 10]. These include organoids [11, 12], where cells are allowed to organize in three dimensions (3D), or organ-on-a-chip devices [13], where the microfluidic device represents the organ geometry and the microfluidics enable temporal control of the flows and physical conditions. Recent work has also dealt with combining the advantages of both approaches to produce organoids-on-a-chip [14]. Gathering general rules from these systems that would explain the outcome of a CTL attack remains challenging as it requires to link quantitative measurements of T cell behavior, which is inherently stochastic, with tumor cell fate [9, 14].

Here, we introduce a microfluidic-based approach for the multiplexed analysis of tumor spheroid fate in the presence of defined immune cell populations. This methodology is based on the parallel formation, manipulation and observation of hundreds of tumor spheroids within stationary microfluidic droplets. This is combined with a dedicated image analysis pipeline to extract immune cell dynamics and tumor spheroid fate within individual droplets [15, 16]. When associated with mathematical models, the quantity and quality of spatiotemporally resolved data allow us to pinpoint key behaviors leading to spheroid destruction and to detect and understand heterogeneity of tumor outcomes. The resulting *Multiscale Immuno-Onclogy on-Chip System* (MIOCS) is applied to understand CTL attack on melanoma spheroids, allowing us to establish the importance of collective CTL behaviors in promoting tumor regression.

## II. RESULTS

### A. Parallelized immune challenge on an integrated microfluidic chip

The immunogenic rejection of 3D cancer models is studied using the classic model of Ovalbumin-expressing mouse B16 melanoma (B16-OVA), which are challenged by OVA-specific CD8+ cytotoxic T lymphocytes (CTLs) bearing the OT-I transgenic TCR (see methods). The complete experiments rely on a microfluidic device that consists of a droplet generating region followed by a droplet trapping region that serves to culture the cells and observe them (Fig. 1a,b) [15, 17]. This trapping region is patterned with 234 microfluidic anchors [18], that allow the droplets to be held in place even in the presence of an external flow. The anchors are diamond-shaped (see Fig. 1b) in order to allow for multiple droplet pairings [16].

**Figure 1.**
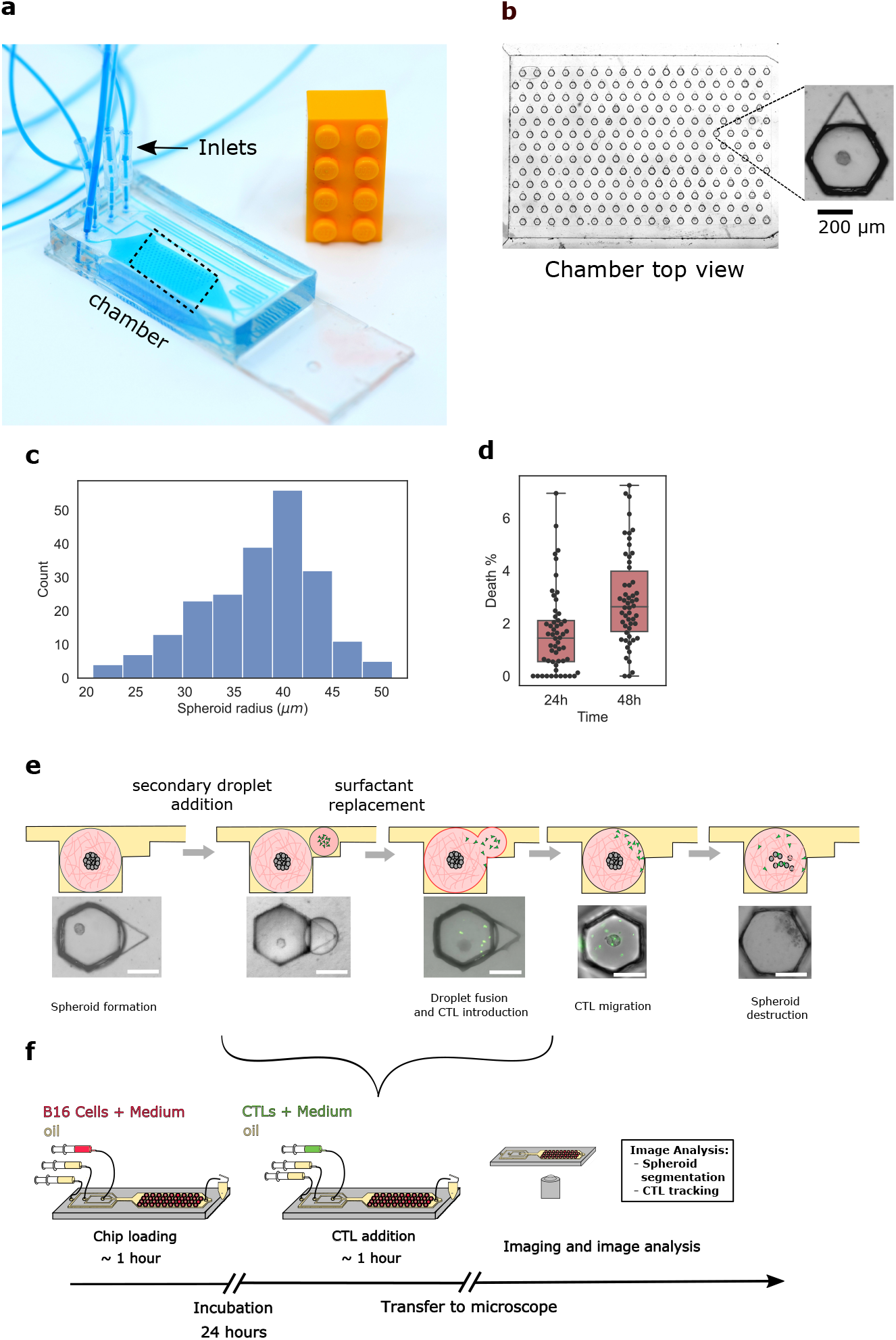
Microfluidic immuno-oncology chip and protocol. **a** Microfluidic chip on a standard glass slide. **b** Expanded view of the trapping region of the chip (dashed box) showing an array of 234 trapped droplets. Each droplet contains a single B16 spheroid in Matrigel, as shown in the inset. **c** Distribution of spheroid radii within a single chip (n=215). **d** Viability measurements using live-dead staining after 24h and 48h (n = 54). **e** Schematic showing a primary droplet with a tumor spheroid, followed by the addition and fusion of a secondary droplet containing GFP labeled CTLs, eventually leading to tumor cell killing and spheroid fragmentation. Scale bar is 200*μ*m. **f** Schematic representation of the complete experimental protocol.

The experiment begins by producing aqueous droplets (Volume = 50 nl) containing Matrigel and a suspension of B16 cells at a concentration of 1.5 · 10^6^ cell/ml. To obtain single spheroids in each droplet, a concentration of 2.0 mg/ml of Matrigel was used, with higher Matrigel concentrations leading to the formation of multiple spheroids per droplet (Fig. S1). Once the droplets are anchored, the device is placed in an incubator at 37°C overnight, which allows a single B16 spheroid to form in each droplet (Fig. 1b). At this cell and Matrigel concentrations we obtain spheroid radii in most cases ranging between 35 to 45 *μ*m (Fig. 1c). A live-dead staining shows that less than 3% of the cells were dead after 48h in the chip (Fig. 1d).

After overnight incubation the CTLs are brought to the Matrigel droplets by generating a group of secondary smaller droplets (Volume = 10 nL) that contain a Poisson distribution of CTLs (Fig. S1b). These secondary droplets are trapped in the triangular sections of the anchors and then merged with the spheroid-containing Matrigel, thus bringing the two cell populations into the same Matrigel droplet (Fig. 1e) [16]. The interactions between the CTLs and spheroids are observed by time-lapse imaging, typically over 24 hours. The whole process from spheroid preparation to CTL addition and imaging is done on a single chip (Fig. 1f) and each experiment yields up to 234 individual replicates, of which we typically obtain 50 time lapse movies, due to the small image acquisition time-intervals (2min/frame). For higher time-intervals, more data points can be collected from the same chip.

Three stages in the cell-cell interactions can be identified: the CTL exploration of the gel, their accumulation on the spheroid, and the killing of B16 Ova-expressing cells (see **Movie 1** for representative cases). These stages are studied in detail below.

### B. CTL migration in micro-device reproduces *in-vivo* behavior

The CTL migration is tracked in the time lapse movies as the cells perform 3D migration within the gel or on the surface of the spheroid (see Fig. 2a). The recorded velocities display alternating periods of motility and arrest phases, as seen by the high and low velocities in Fig. 2b. This behavior, as well as the value of the velocities, correspond well to previously reported CTL velocities in collagen gels *in-vitro* [19] or within tissues *in-vivo* [20, 21]. We can infer the 3D motility properties from the acquired microscopy data as shown in SI section VI A. Note that the value of the measured velocities depend on the image acquisition frequency (see SI section VI B and Fig. S2a,b) In order to evaluate the influence of the spheroid presence in the droplet on the motility of the CTLs, the migration statistics without a spheroid present are compared with the statistics in the presence of a spheroid during the first 500 minutes of an experiment. We restrict the analysis to cells that are not in contact with the spheroid. The displacement distributions (Fig. 2c) and the mean-square displacements (MSD) (Fig. 2d) do not show any significant difference between the two conditions. In both cases, the CTLs undergo super-diffusive random walks (Fig. 2d) with MSD ~ *τ ^α^* and *α* = 1.6, in agreement with what was reported *in-vitro* and *in-vivo* [19, 21].

**Figure 2.**
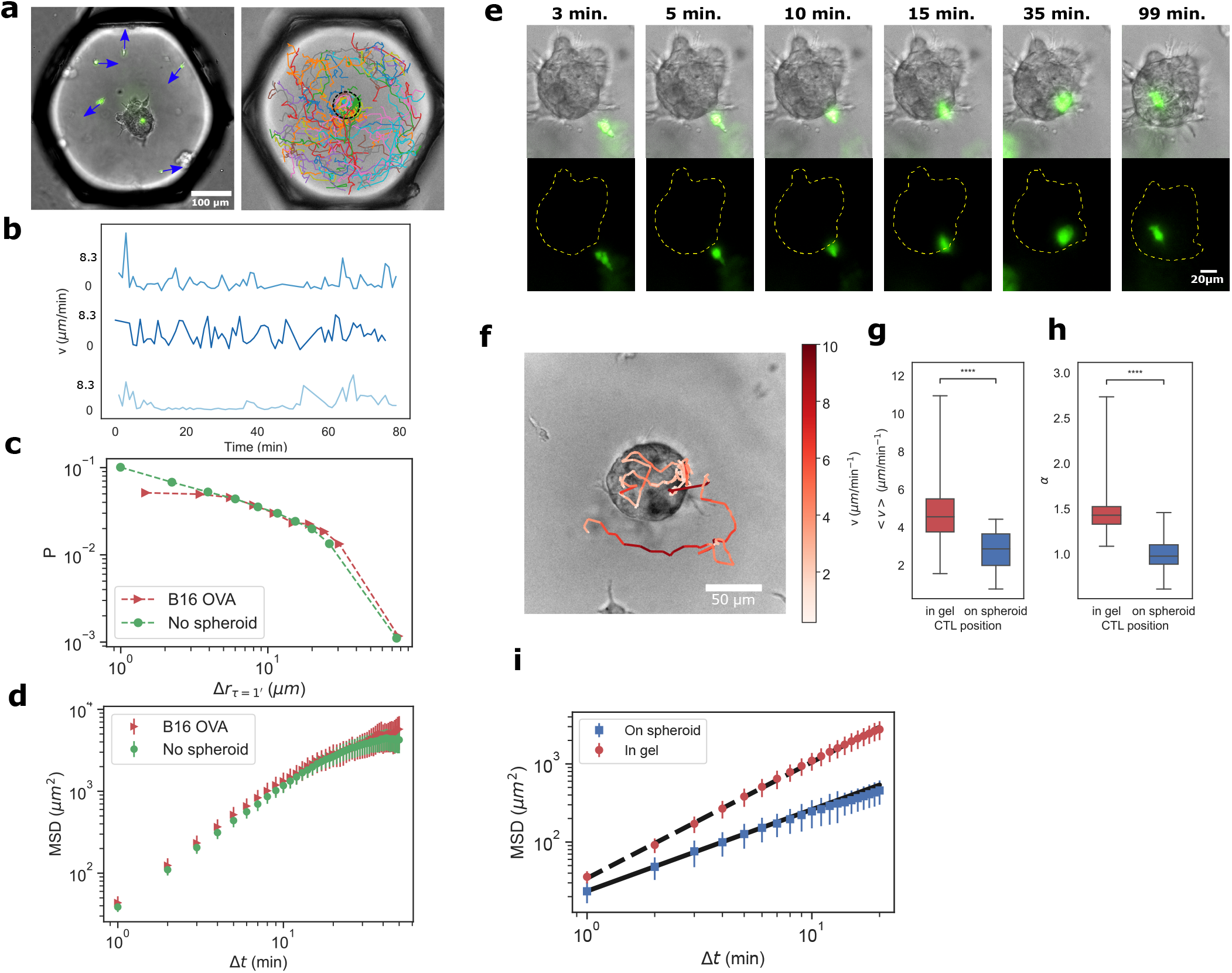
CTL migration in droplets recapitulates *in-vivo* behavior. **a** (left) Representative image of CTLs with instantaneous velocity vectors inside Matrigel droplet. (right) CTL tracks in one droplet over 24 h, each color represents an individual cell track. The dashed black circle outlines the spheroid boundary. **b** Representative velocities as a function of time for three different T-cells. **c** Probability distribution of a cell to migrate by a given distance (∆*r*) during a fixed time step ∆*t* = 1 minute (*n* = 67965 points without spheroid and *n* = 34072 individual points for CTLs in presence of the B16 spheroids). **d** Mean Square Displacement (MSD) of CTL migration with and without spheroids. **e** Time sequence showing the initial CTL approach and contact with a spheroid. **f** Track of a single CTL as it migrates in the matrigel and on the spheroid surface. Colormap represents the instantaneous velocity of the cell. **g, h** Average velocity and mean square displacement exponent (*α*) of cells migrating in the gel and on the spheroid. Each data point is the average velocity in a given droplet (*n*_gel_ = 55, *n*_spheroid_ = 54). **i** Mean-square displacement of cells migrating in the matrigel and on the spheroid. Bold and dashed lines represent the best fits for the MSD of CTLs on the spheroid and in the matrigel, with respective exponents of 1.1 and 1.4.

After some time, one CTL comes in contact with the spheroid. This contact generally leads the T-cell to adhere to the spheroid and explore its surface over the course of a few hours (Fig. 2e,f and **Movie 2**). We select individual tracks with segments both on and off the spheroid to investigate more precisely the CTL motility change upon reaching the spheroid. We observe that the CTL behavior is strongly modified: They display lower mean velocity (Fig. 2g and **Movie 2**) with a lower MSD exponent ( Fig. 2h,i). The average MSD exponent goes from 1.4 when the cells move in the gel to 1.1 after the same cells have reached the spheroid.

The CTL migration in the gel therefor recapitulates behaviors that have been reported *in-vivo* [19–22], with the current data highlighting the switch in motility before and after the CTL contacts the spheroid surface.

### C. A positive feedback loop drives CTL accumulation on the spheroid

We now investigate the contact time statistics of the CTLs on the spheroids. The distribution of first-contact times is consistent with the distribution of randomly migrating particles in an enclosed environment (see Section VI E [23]), further indicating that the initial contact is random and that there is no attraction from the spheroid on the CTLs (Fig. 3a). After the first CTL contact with the B16 cells, the arrival of successive CTLs leads to an accumulation of T cells on the spheroid in the case of the B16-OVA spheroids. This accumulation is shown in absolute numbers (Fig. 3b) and also by computing the ratio of CTLs in each droplet that are present on the spheroid as a function of time (Fig. 3c). However the accumulation is not observed in the case of wild-type B16 cells, which do not express Ova (**Movie 3**).

**Figure 3.**
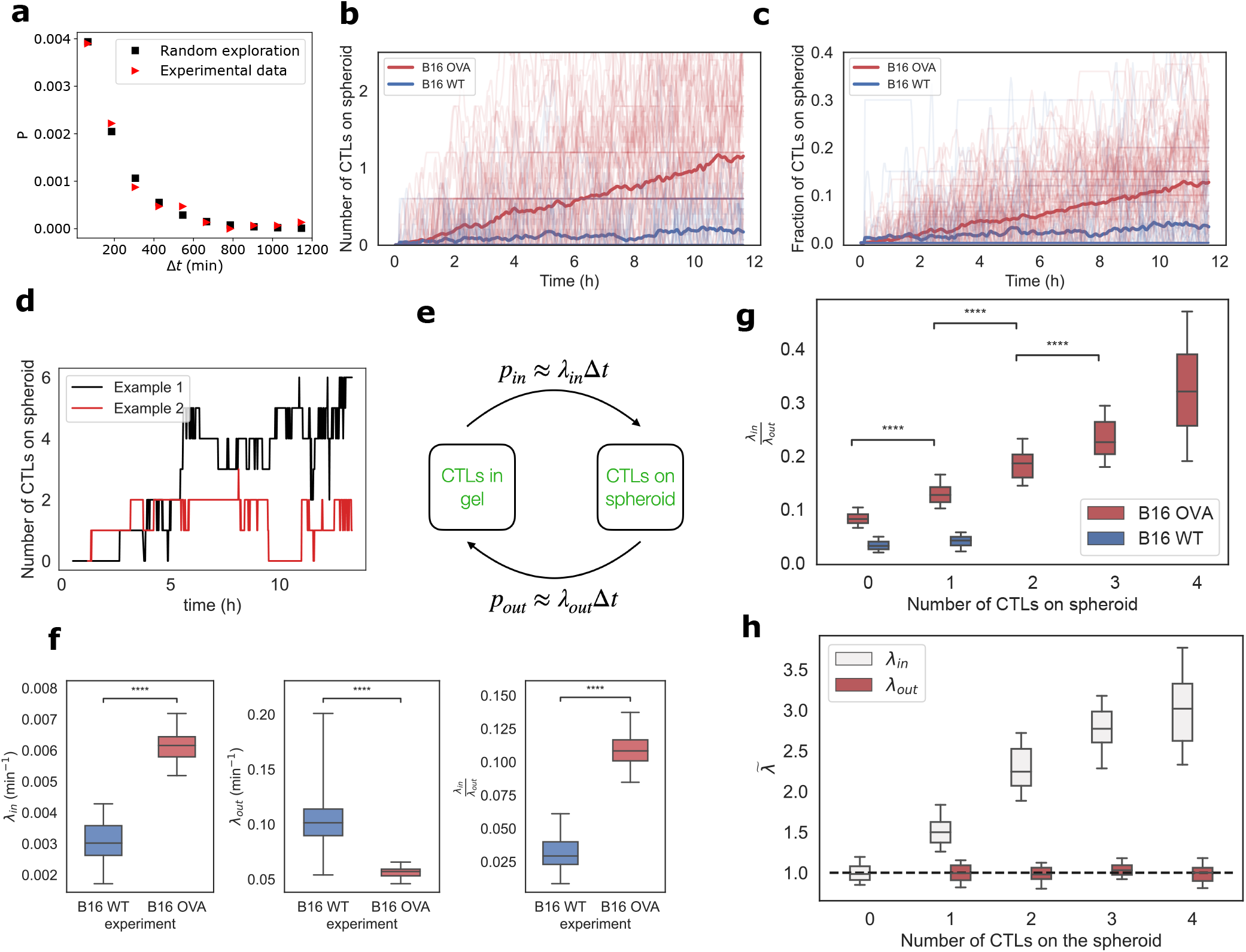
CTL accumulation enhanced by a positive feedback-loop after first contact. **a** Experimental distribution of first CTL-spheroid contact times and theoretical distribution for randomly migrating CTLs. **b** Number of CTLs detected on each spheroid and **c** the fraction per droplet as a function of time. Each thin line represents a single tracked spheroid, in bold is the averaged value. In red is the accumulation for B16-OVA spheroids and in blue for B16 WT spheroids. [84 individual B16-OVA spheroids and 81 B16 WT spheroids tracked] **d** Number of CTLs as a function of time on two representative spheroids showing the detection of attachment/detachment events. **e** Schematic of the stochastic accumulation model: CTLs can switch from the gel to the spheroid with different probabilities. *p_in_*(∆*t*) (conv. *p_out_*(∆*t*)) is the probability for a cell to attach to (conv. detach from) the spheroid during a time interval ∆*t*. Counting attachment and detachment events in the experiments allows us to infer the rates *λ_in_* and *λ_out_*, see section VI E for details. **f** Estimates for the attachment rates (*λ_in_*), detachment rates (*λ_out_*) and affinity ratio (*λ_in_ / λ_out_*) for B16-WT (blue) and B16-OVA (red) cells. The box plots are obtained by using a bootstrapping method with 50 repetitions as described in the methods. **g** The affinity ratio as a function of the number of CTLs attached to the spheroid for B16 WT and B16 Ova spheroids. **h** Normalized attachment rate *λ_in_* (white) and detachment rate *λ_out_* (red) as a function of the number of CTLs attached to the spheroid. *λ_in_* and *λ_out_* are normalized by their mean values for 0 and 1 CTL on the spheroid respectively.

At this stage an important question is whether this accumulation results from cells reaching the spheroid randomly, as they explore the droplet volume, or if the accumulation rate is enhanced due to cell-cell signaling. We address this question by analyzing the accumulation of the CTLs at the spheroid level. At this scale, the CTL accumulation is not homogeneous but stochastic, with CTLs both attaching and detaching over time (Fig. 3d, **Movie 2**, **Movie 4**). The measured attachment and detachment statistics allow us to compute the cell attachment rate, *λ_in_*, as well as the detachment rate, *λ_out_* (Fig. 3e), each of which is computed on the spheroidlevel in each experiment, as explained in the SI (see Sec. VI E).

The value of *λ_in_* is found to be significantly higher in the B16-OVA spheroids compared the B16-WT spheroids, while the opposite is true for *λ_out_* (Fig. 3f). These measurements indicate that the arrival rate of CTLs increases when the cells composing the spheroid express the cognate antigen recognized by the CTLs and that they stay attached for longer periods of time. Therefore the accumulation of CTLs on the spheroids is mediated by two independent phenomena, first the increase in arrival frequency and second by the decreased leaving frequency. The net effect of the attachment/detachment dynamics can then be summarized by the affinity ratio, *λ_in_ / λ_out_*, which accounts for the net effective accumulation of CTLs on target. This ratio is found to be significantly higher with B16-OVA spheroids when compared to the B16-WT spheroids (Fig. 3f).

Evidence of a positive feedback loop for the attraction among the CTLs can be obtained by calculating the change of the affinity ratio as a function of the number of cells present on the spheroid. Indeed, the depth of the experimental data allows us to obtain a value of *λ_in_ / λ_out_* before the first contact and then successively track the change of this ratio after every contact, in each droplet (Fig. 3g). Again, the data for B16-OVA show a significant difference with the WT case. More interestingly, the higher the number of CTLs present on the B16-OVA spheroid, the higher the ratio, and thereby, the faster the accumulation rate: this is a hallmark of a positive feedback. The increase in the accumulation rate is driven by an increase in *λ_in_* for a constant value of *λ_out_* (Fig. 3h). It demonstrates that the accelerated accumulation is mediated by the increasing attraction of the CTLs to the spheroid, which is triggered from the very first CTL attached to the spheroid (Fig. 3h).

The current results confirm recently published observations [24] that show a CCR5-mediated swarming of T-cells *in-vitro*, as well as *in-vivo* studies that report the accumulation of T-cells on targets [25, 26], Figure 3 shows that these effects occur even for cell populations consisting of a few individuals and that a single contact can trigger the beginning of the positive feedback. Below we go beyond the CTL accumulation to address the relationship between the accumulation of CTLs and their capacity to kill the B16 spheroids.

### D. Killing of B16 cells by CTLs is heterogeneous

After focusing on the behavior of the T-cells we now turn to the response of the cancer spheroids upon CTL accumulation. The time-lapse movies allow us to identify individual cell death events in the spheroids at the molecular level by detecting the activation of Caspase 3/7, which provides an early marker of apoptosis [27] (Fig. 4a and **Movie 5**). In the bright-field image we observe instances of rapid shedding of cellular material and debris from the spheroids (Fig. 4a, **Movie 1, 3, 5**), which we refer to as "spheroid fragmentation". Combining this information with the position of CTLs relative to the spheroid, it is thus possible to record a detailed chronology of the key events taking place in each droplet by timing successive CTL contacts with the spheroid and the apparition of fragmentation events and Caspase 3/7 signals, as shown in Fig. 4b.

**Figure 4.**
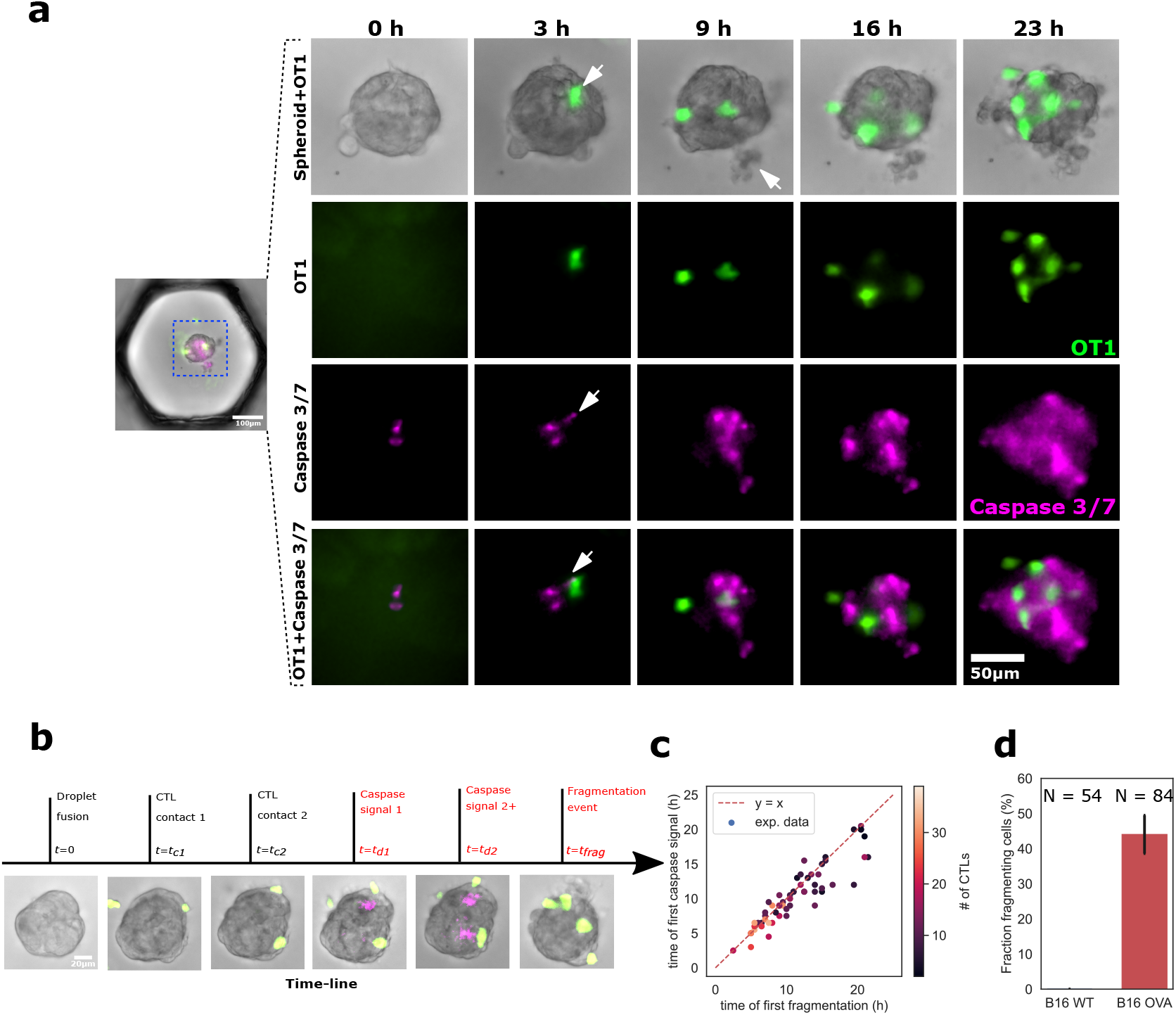
CTLs kill tumor cells within spheroids. **a** Representative sequence showing CTL positions on the spheroid, caspase 3/7 fluorescent death marker, and B16 cell fragmentation. The white arrow at 3h indicates the appearance of capsase signal next to a CTL. At 9h it represents a fragmented dead cell. **b** Representative chronology showing the key events for a given spheroid interaction with CTLs: Contact times of CTLs on spheroids, detection of caspase 3/7 signal, detection of fragmentation events. **c** Time of first caspase signal vs. first observation of cell fragmentation. **d** Percentage of WT (black) and OVA (red) spheroids that show at least one fragmentation event in under 14 h. *N* equals 54 and 84 spheroids respectively.

We observe that the timing of the first Caspase event post-CTL contact is well correlated with the first fragmentation event (Fig. 4c), indicating that the two observations are closely related. For this reason, we will hereafter use brightfield images to quantify spheroid killing, which simplifies the analysis pipeline. Furthermore the timing of these killing events is highly variable, ranging from a few hours to beyond 24 hours. For some spheroids no fragmentation or Caspase events are observed over the course of an experiment. In the following analysis we will label “successful killing” the cases when the first fragmentation event is observed before t=14 h. The statistics of such events are summarized in Fig. 4d, which shows that 44% (*N_OV A_* = 84 spheroids) of the OVA expressing spheroids are successfully killed by the CTLs. This contrasts with the B16-WT spheroids, where we do not observe any fragmentation events (Fig. 4d, **Movie 3**). An analysis of the statistical impact of each of the problem parameters on a successful killing shows that the dominant parameters are the number of CTLs in the droplet and the number of CTLs that reach the spheroid (Section VI C).

### E. Tumor spheroid killing involves collective effects

We now consider the relationship between the spatiotemporal dynamics of the CTLs and the tumor spheroid outcomes (successful or unsuccessful killing). An indication of the relevance of this link is obtained first by observing that the CTL accumulation rates are faster on the spheroids that display fragmentation than in the opposite case, both in absolute numbers (Fig. 5a) and as a fraction of the total number of cells per droplet (Fig. 5b). This indicates that a faster accumulation is correlated with efficient killing.

**Figure 5.**
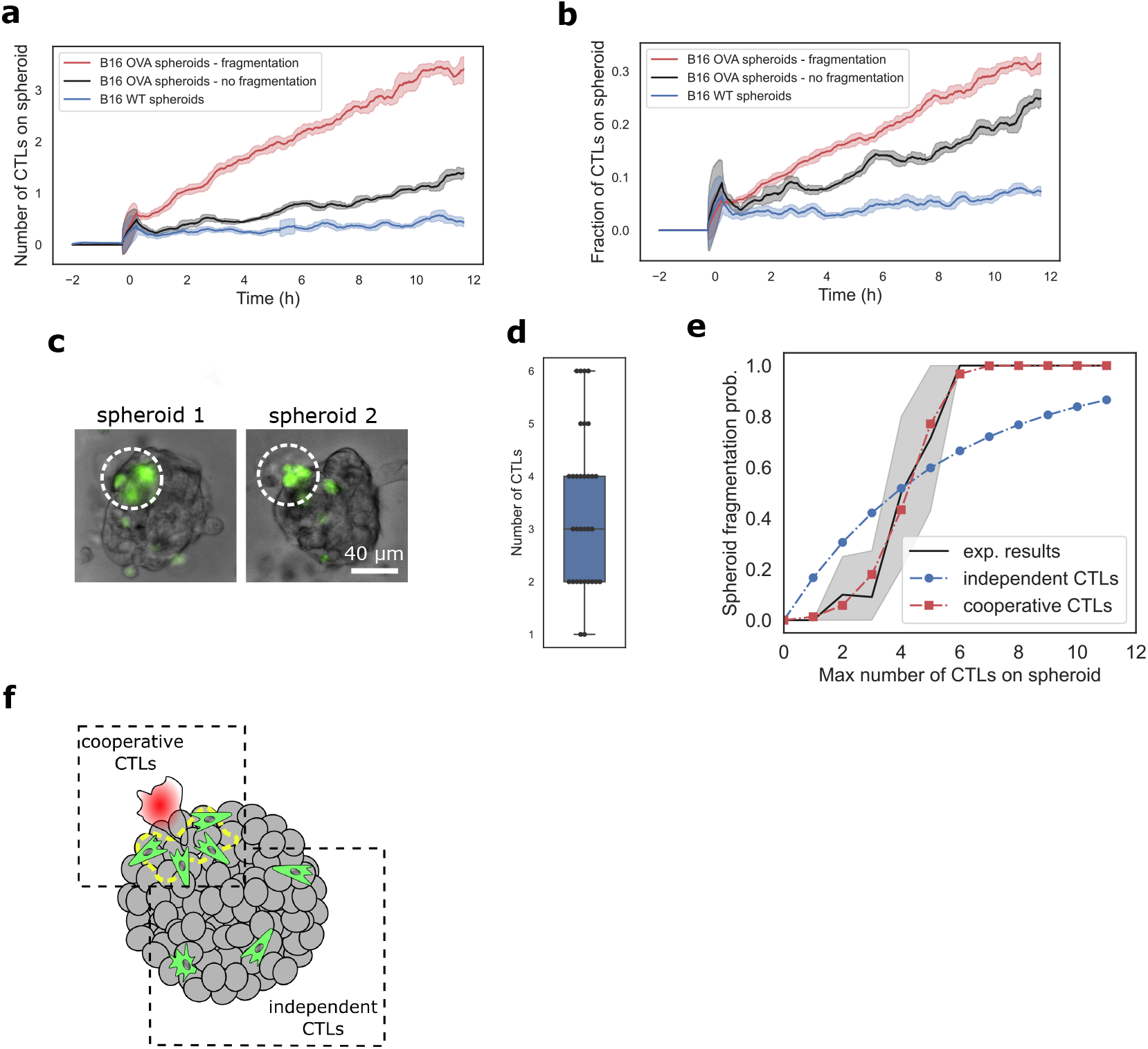
CTL number and collective behavior determine probability of killing. **a, b** Number (a) and fraction per droplet (b) of CTLs on fragmenting (red) and non-fragmenting (black) spheroids as a function of time (re-centered according to the first contact time). **c** Two representative images showing CTL clustering on the spheroid during first fragmentation event. The white circles have a radius of 30*μm* around fragmenting cell. **d** Distribution of the CTL numbers within a radius ≤ 30*μm* around the fragmentation areas (*N* = 31). **e** Probability of spheroid fragmentation as a function of the maximum number of CTLs observed on the spheroid. Experimental measurements (*N* = 84, black line) are compared with the results of the independent CTL model (blue dots), cooperative model (red squares). Shaded area gives the 95% confidence interval of experimental data. **f** Sketch summarizing the observed behavior: CTL (green) clustering leads to tumor cell killing and fragmentation (red).

Moreover observing the CTLs on the spheroid reveals that fragmentation events are associated with the presence of several CTLs in the vicinity (Fig. 5c,d). This local effect is quantified by counting the number of T cells present in within a 30 *μ*m radius of the first cell fragmentation event, giving a mean number of 3.4 cells (median at 3 cells), with fragmentation very rarely occurring with only one cell present at the tumor site (Fig. 5d). Indeed, CTLs sometimes appear to besiege a salient B16 cell, causing it to burst after a few minutes of attack.

While the above observations suggest the possibility of a cumulative effect of the CTLs that enables the killing of tumor cells, they do not directly assess cooperation where the CTLs interact together to achieve the killing. The existence of this cooperative killing is evidenced by measuring the probability of a spheroid to display fragmentation as a function of the maximum number of CTLs present on it over the course of the experiment (Fig. 5e). The experimental measurements are compared with an independent and a cooperative probabilistic model (for more details see SI section VI D): For the independent killing model, we hypothesize that each contact between a CTL and a target cell has a given probability to cause a fragmentation event on the spheroid. This hypothesis leads to the parabolic probability of killing, as shown by the blue dots on Fig. 5e. The poor match with the experimental data demonstrates that a model of independent interactions between the CTLs and the target cells fails to capture the underlying mechanism.

Instead, a much better fit can be obtained by supposing that the probability of a CTL to lead to a fragmentation event depends on the number of other CTLs present on the spheroid. As shown in Fig. 5e,a simple model of cooperative interaction leads to a distribution that closely matches the experimental data, thus providing further support for the cooperative aspect of the killing. Taken together the elements of Fig. 5a-e indicate that the CTLs tend to cluster at particular sites on the spheroid and that their clustering enhances their ability to induce spheroid fragmentation (Fig. 5f).

### F. Long and short-range interactions combine to determine probability of CTLs to kill a tumor spheroid

The probability for successful killing to occur for any particular spheroid can now be explained as a combination of the effect of CTL accumulation on the spheroid and their collaborative killing behavior. This probability is a function of the total number of CTLs in the vicinity of the spheroid, which also correlates with the maximum number observed on the spheroid (Fig 6a). It can be modeled using the parameters computed above for the two collective behaviors (Fig 6b).

**Figure 6.**
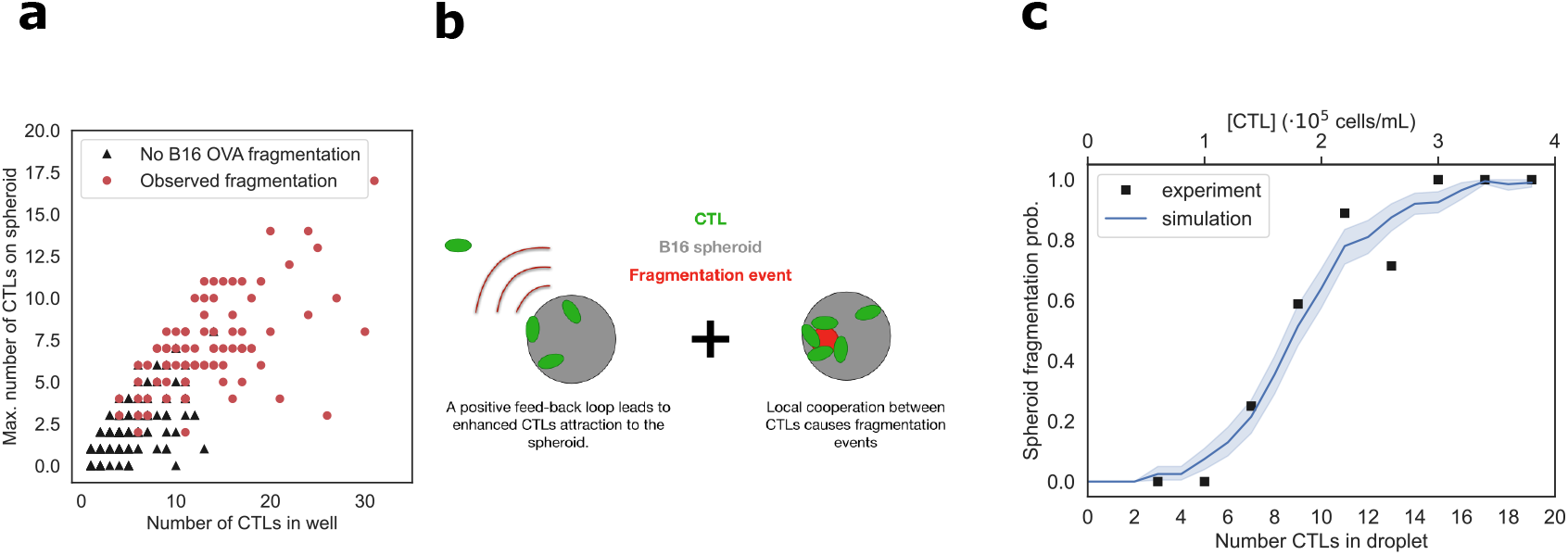
Combining short and long-range interactions to predict spheroid fate. **a** Spheroid fragmentation as a function of the number of CTLs in the droplet and the maximum number of CTLs detected on the spheroid. **b** The concentrationdependent killing is modeled as the result of two complementary mechanisms: long-distance cooperative attraction of CTLs to the target site and local killing cooperation on the spheroid. **c** Experimental (black squares) and simulated (blue line) spheroid fragmentation probability as a function of the number of CTLs in the droplet. Shaded area represents the 95% confidence interval of simulated data.

For a given number of CTLs in the droplet, the accumulation of CTLs on the spheroid is simulated using the estimated hitting and leaving probabilities described in Fig. 3. The final state of the spheroid (fragmented or not fragmented) is then drawn using a Bernoulli distribution whose parameter depends on the maximum number of CTLs on the spheroid using the probabilities of Fig. 5e. This computation is performed for a number of CTLs per droplet ranging from 0 to 20 cells, repeated 200 times. The average outcome yields the simulated fragmentation probability curve as a function of the number of CTLs in the droplet (Fig. 6c). Not only does the simulation confirm the key role of CTL concentration in initiating spheroid fragmentation, the close match between simulated and experimental measurements (with no free parameters) indicates that the spheroid fragmentation process is well described from this succession of two mechanisms: cooperative recruitment and collective killing at the spheroid site.

## III. DISCUSSION

The current study introduces a new paradigm for extracting biological information from *in-vitro* experiments, by treating the parallel realizations as *“Monte-Carlo experiments”* that reach different outcomes in a probabilistic way. This contrasts with existing microfluidic models for cancer-immune interactions, which treat each chip as a single experiment and use traditional biological measurement techniques [10, 28]. By comparison the droplet format provides several unique features, including the ability to merge many droplet pairs at a well-defined time, thus providing a common starting time of the parallel experiments [16, 29]. Moreover, the encapsulation within droplets allows the conditions in each of the parallel experiments to be well controlled, thus allowing for massively multiplexed experiments on a single device.

Here these technical advantages are associated with probabilistic modeling to infer key biological information about the ability of CTLs to sense and respond to the tumor, by relating the spatiotemporal dynamics of the CTLs with the outcome for the tumor spheroid. Specifically, we find that the first CTL-tumor cell contact, which occurs randomly, triggers a positive feedback loop that leads to an accelerated accumulation of CTLs on the spheroid. Later, CTLs form clusters on the spheroid that enhance their ability to kill the target cells, leading to tumor rejection.

Several mechanisms may account for the collaborative CTL accumulation and killing. Chemokines that are both sensed and produced by T cells have the ability to drive their swarming behaviour [24]. Cooperative killing, in which multiple sublethal cytotoxic hits synergize to induce target cell killing, has been previously described in the context of viral infection [30] and tumor development [31]. Alternatively, initial CTL-tumor cell interactions may facilitate tumor destruction by increasing MHC class I expression through IFN-*γ* production and diffusion in the tumor microenvironment for example [20, 32]. As illustrated here, our approach helps support and generate new hypotheses that can be subsequently dissected at the molecular level.

Looking ahead, the Multiscale Immuno-Oncology onChip System (MIOCS) can now be generalized to include several immune cell types and more realistic tumor models in each droplet. Here again the spatiotemporal resolution and *Monte-Carlo* approach will be fundamental to understand the causality of the interactions and the effect of 3D geometry. Finally, working with patient-derived organoids [33, 34] will have important implications for personalized medicine.

## ACKNOWLEDGEMENTS

The authors acknowledge enlightening discussions with Clément Roussel, Andrey Aristov, Raphael Tomasi, Gabriel Amselem and all members of the Baroud team. PB is funded by Institut Pasteur, Inserm and an advanced grant from the European Research Council (ENLIGHTEN). GR is funded by the Direction Générale de l’Armement. The authors acknowledge the support of the Biomaterials and Microfluidics platform and the FabLab at Institut Pasteur.

## AUTHOR CONTRIBUTIONS

GR, SJ, MC, PB, CNB designed experiments. GR, SJ performed experiments and analysis. CA, MC and RK performed experiments. GR developed mathematical models, image and data analysis tools. PB and CNB supervised the research. GR, SJ, PB, CNB wrote the manuscript. All authors discussed the results and commented on the manuscript.

## IV. EXTENDED DATA FIGURES

**Figure S1.**
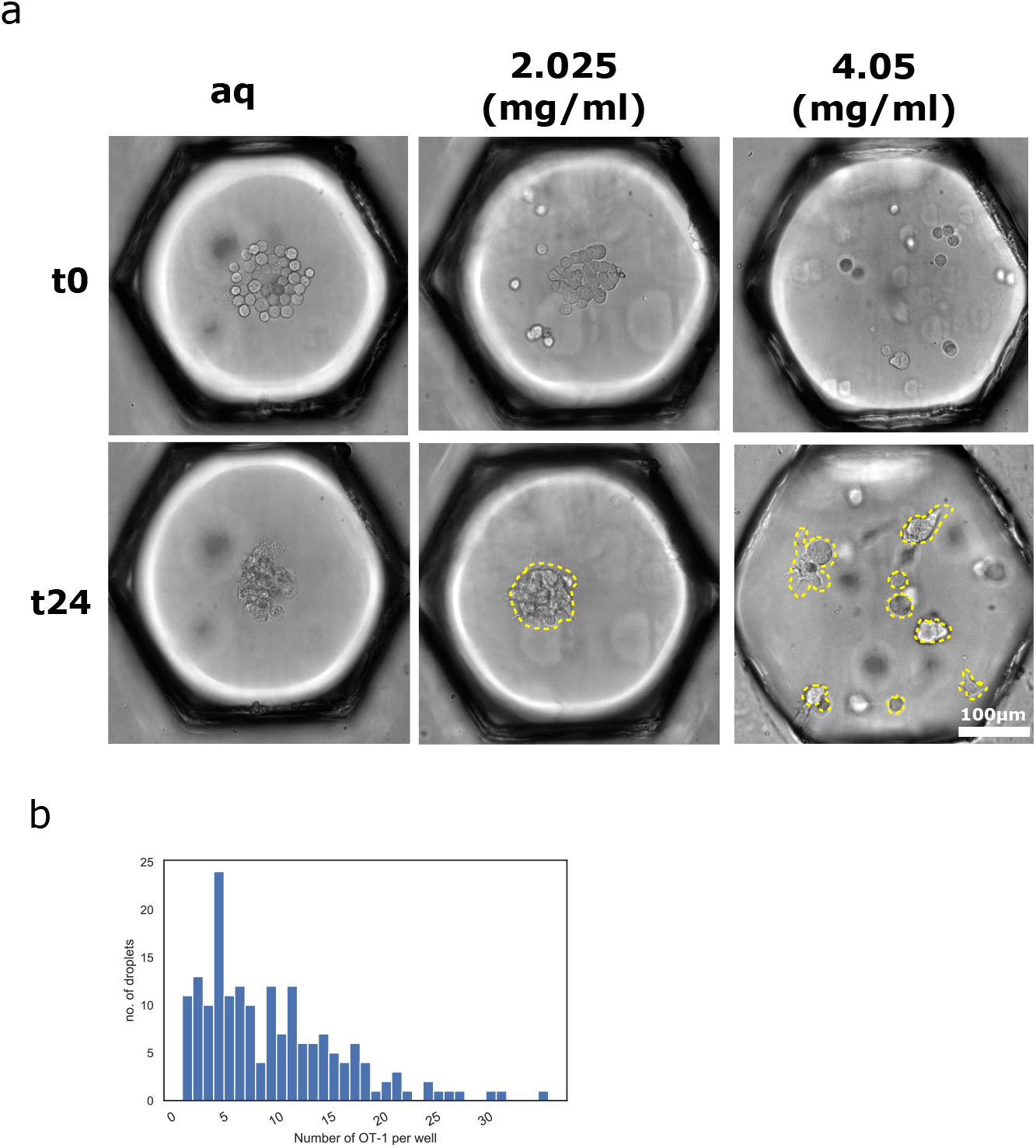
Extended data fig. 1. **a** Representative spheroids for different Matrigel concentrations at *t*_0_ and *t*_0_ + 24*h* **b** Distribution of the number of CTLs per droplet (*n* = 151)

**Figure S2.**
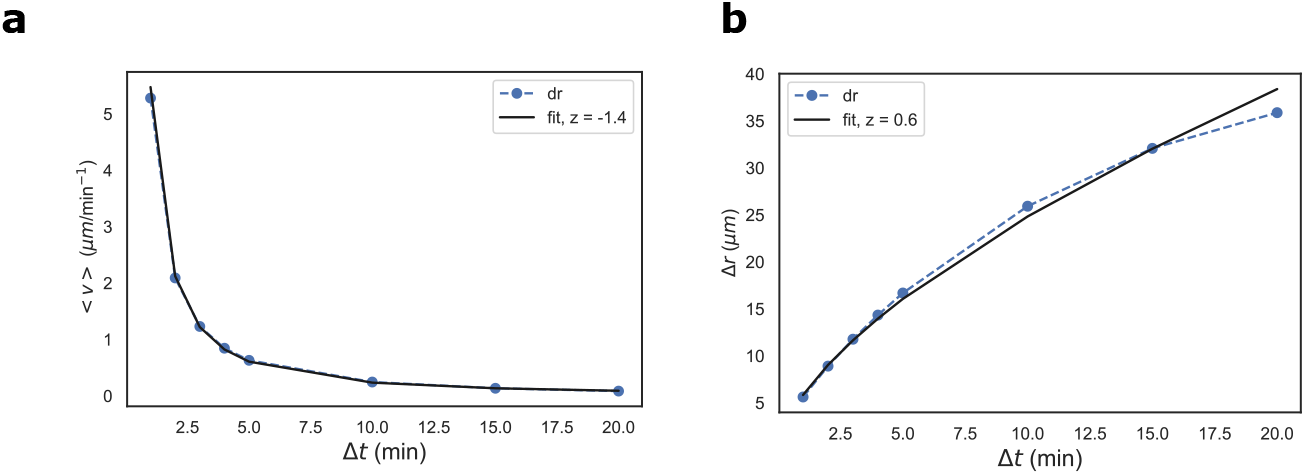
Extended data fig. 2. **a** Measured velocity as a function of the time-interval between two successive position measurements. The velocity scales as a power law. **b** Measured traveled distance as a function of the time-interval between two successive position measurements.

## V. MATERIALS & METHODS

### a. tumor cells

B16-WT melanoma and B16OVAlbumin peptide (residue 257-264) expressing celllines (B16-OVA) were maintained in RPMI 1640 (Fischer Scientific – 12027599) media containing 10% FBS and 1% penicillin-streptomycin antibiotics and were maintained at 5% CO2 and 37°C. B16.F0 (B16) and B16.F0Ova (B16-OVA) melanoma cells were kindly provided by Claude Leclerc.

### b. Generation of OVA-specific cytotoxic T lymphocytes (CTLs)

Ubi-GFP Rag1l/ OT-I TCR mice were bred in our animal facility under specific pathogen–free conditions. Splenocytes were isolated from Ubi-GFP OTI TCR transgenic mice and red blood cells were removed by ammonium–chloride–potassium lysis. One-third of the cells was then pulsed with 50 *μ*M of Ova257–264 peptide (SIINFEKL) for 2h at 37°C in 1 mL total volume of RPMI medium 1640-GlutaMAX. The rest of the cells were incubated at 37°C in 15mL of complete medium (RPMI medium 1640-GlutaMAX supplemented with 10% heat-inactivated fetal bovine serum, 50 *μ*g/mL penicillin, 50 *μ*g/mL streptomycin, 1 mM sodium pyruvate, 10 mM HEPES and 50*μ*M *β*-mercaptoethanol). After 2 hours, the two populations were mixed and cultured for 3 days. Cells were then subjected to Ficoll gradient centrifugation to remove dead cells and thus select live Ova-specific CTLs, and cultured in complete RPMI medium, supplemented with human IL-2 (10 ng/mL; R&D) for 2 additional days.

### c. Microfabrication, Microfluidic setup and Droplet formation

The PDMS-based microfluidic device on which the experiments were conducted is precisely described in Refs. [15, 16, 35]. Preceding droplet production, the chip in filled with fluorinated FC40 (3M) oil mixed with 2%(v/v) FluoroSurfactant (Ran Biotechnologies) and cooled at −20°Cfor two hours to prevent Matrigel gelification during the loading. The primary droplets are produced as in Sart et al. [15]. The aquous phase is composed of RPMI media, Corning Matrigel (Dutscher Dominique – 354234) and B16 melanoma cells at a concentration of 1 · 10^6^ cells/mL. The primary droplets have a volume of ca 50nL. The chip is then placed in the incubator at 37°Cleading to Matrigel gelification.

The secondary droplets (volume ca. 10nL) are produced with an aqueous phase containing RPMI media and OT-1-CD8+GFP cells at a concentration of 1 · 10^6^ cells/mL. The secondary droplets were further trapped in the triangular regions adjacent to the individual hexagonal wells already present with primary droplets encapsulated with B16 spheroids (see fig. 1). To fuse the primary with the secondary droplets, 20%(v/v) of 1H,1H,2H,2Hperfluoro-1-octanol (PFO) (Sigma-Aldrich) was dissolved in NovecTM-7500 Engineered Fluid (3M) and was perfused in the microfluidic chip. This causes the fusion of adjascent droplets. After the fusion of droplets, a fresh solution of FC40 and Fluorosurfactant was flushed-in to remove the PFO in the microfluidic chamber.

### d. Spheroid formation in different matrigel concentrations

We primarily used the Matrigel concentration of 2.0 mg/ml in order to encapsulate the droplets with B16 cells. 24 hours after droplet loading, the B16 cells self-assemble into a single 3D spheroid of B16 tumor cells (Fig. 1b). Using concentrations higher than 2.0 mg/ml (4.05 mg/ml) often leads to multiple spheroids (Extended Data Fig. S1a) located at different droplet heights. Conversely droplets with pure aqueous media (without Matrigel) resulted in single spheroid formation but lacked the Extra-cellular Matrix (ECM) necessary for the migration of T-cells (Extended Data Fig. S1a).

### e. Viability assay (Fig. 1)

B16-OVA spheroid containing droplets were made according to the protocol described above with the addition of Propidium Iodide (PI) (Sigma P4864) at a concentration of 3*μ*M. The chip was then imaged at 24 and 48 hours after seeding. Only spheroids positioned in the center of the droplets were imaged, in order to avoid artifacts due to the microscopy. Using a custom-made Imagej macro, the area of red fluorescent signal by PI was measured and the percentage fluorescent area was calculated when compared to the complete spheroid area.

### f. Apoptosis assay (Fig. 4)

Caspase-3/7 Red Apoptosis Assay Reagent (Essen bioscience – 4704) was used at 2*μ*M concentration (added during primary droplet formation) in order to visualize the apoptotic cells in the spheroids.

### g. Microscope imaging

Images were captured using a Nikon Ti2 motorized epifluorescence microscope with a 20x objective lens. The illumination was produced by a Lumencor LED light source and the images were captured by a Hamamatsu C13440-20CU SCMOS camera.

### h. Image analysis

A specific image analysis pipeline was developed in order to extract physical and biological variables from the time-lapse movies. The routines were written in Python programming language and use several open-source libraries [36–38]. The code is available upon reasonable request.

### i. Determining the positions of CTLs with regard to the spheroid

Each droplet image is multi-channel, recording a planes in brightfield and the FITC (510 nm) channels. The former enables well identification and spheroid segmentation, whereas the latter enables us to record CTL positions.

To segment the spheroid in a given droplet we rely upon a border-detection routine based on Laplace filtering. The spheroid properties such as size and position are recorded and stored. This procedure is repeated at every time step for more robustness. The CTLs are detected using the fluorescence channel, and their positions are stored. This information is then crossed with the spheroid positions extracted above to determine the position of the CTLs relative to the spheroid in the droplet (on/off the spheroid). For each timestep an image with the raw image, the mask covering the detected spheroid and the relative positions of the CTLs is generated for manual verification a-posteriori of the algorithm efficacy. Faulty wells (with for example mis-detected spheroids) are then thrown away before data analysis. The full image-analysis algorithm is available upon reasonable request.

### j. λ_in_ and λ_out_ estimation and simulation

The attachment and detachment rates are estimated using a bootstrapping procedure. From a subset of the experimental accumulation plots (3d) we count the attachment and detachment events and estimate the values of *λ_in_* and *λ_out_* (see SI VI E for more details). We repeat the procedure 50 times selecting for each iteration 70% of the total number of spheroids in each condition (84 B16-OVA and 81 B16-WT spheroids). This gives us a distribution of the estimated parameters *λ_in_* and *λ_out_* represented in figure 3.

The commented code for estimating the attachment and detachment rates from the experimental data using a bootstrapping procedure are available upon reasonable request. The code used to generate the simulated fragmentation probability curve in Fig. 6c is available at the same address.

### k. Statistical tests

All statistical tests were made using the `statannot` library in Python. Unless explicitely stated otherwise, statistical significance was evaluated using two-sided Mann-Whitney-Wilcoxon tests. The p-value annotation legend is:

- ns: 5 · 10^−2^ < *p* ≤ 1
- *: 1 · 10^−2^ < *p* ≤ 5 · 10^−2^
- **: 1 · 10^−3^ < *p* ≤ 1 · 10^−2^
- ***: 1 · 10^−4^ <*p* ≤ 1 · 10^−3^
- ****: *p* ≤ 1 · 10^−4^

The confidence intervals (Fig. 5e and Fig. 6c) are calculated using the `lineplot` function from the Seaborn plotting library [39], which computes the 95% confidence intervals by bootstrapping the experimental data.

## VI. SUPPLEMENTARY INFORMATION

### A. Interpreting the statistics of cell displacements from 2D images

After the introduction of the T-cells into the Matrigel droplets, they explore the space of the droplet in all three dimensions (3D). For practical considerations however, the imaging was limited to a single plane within the droplet, leading to two-dimensional (2D) slices of the drops. It is Therefore important to consider how to interpret the statistics of 2D measurements of cells moving in 3D. Under the hypothesis of a perfectly isotropic medium, we can study the influence of 2D projection of a 3D motion on the mean square displacement (MSD).

For a displacement during a time-interval *τ*, we can write the following relationship for the position vector **r**:

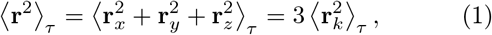

where **r**_*i*_ is the displacement in the direction *i* and 〈·〉 denotes an ensemble average.

The scaling of the mean square displacement as a function of *τ* is not impacted by the projection from 2D to 3D. Indeed the calculation shows that *D* is modified by a numeric coefficient of 2/3. If the motion is isotropic then the relationship between the measurement, which is the projected displacement of the CTL on the x-y plane **r**_*p*_, and the true displacement **r**is:

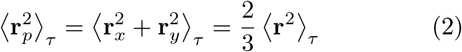

This relationship is useful to assess the link between average projected velocities 〈‖**v**_*p*_‖〉 and average 3D velocities, yet the true relationship is slightly more complicated. Indeed, let us make two hypotheses: the velocity vector **v**is random and isotropic. The angle between the velocity and the imaging plane is *ψ*. Thus:

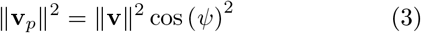

So taking the square root of the velocity vector and averaging over *ψ* we get:

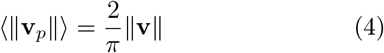

Correcting the velocity values from experimental 2D measurements yield estimated 3D velocities of the order of 7.5*μ*m/min. We have now shown how we can infer, under the conditions of reasonable hypothesis of isotropic cell motility, key properties of 3D cell motility from 2D measurements of the velocity vector.

### B. CTL velocity measurements depend on the imaging frame rate

Measuring the motility properties of particles or cells undergoing random motion is heavily dependent on the sampling rate. Indeed, since the cells can move back and forth, we expect the net distance traveled ∆**r**(*τ*) over a time *τ* to be sub-linear. By measuring the average distance traveled for different lag times we extract the experimental dependence in Fig. S2 b,c. We find that the distance traveled scales as a power law with an exponent smaller than 1 (which corresponds to balistic motion): ∆**r**(*τ*) ∼ *τ*^0.6^. This translates to an experimental average velocity which scales as: **v**(*τ*) ∼ *τ* ^−1.4^.

**Table I.**
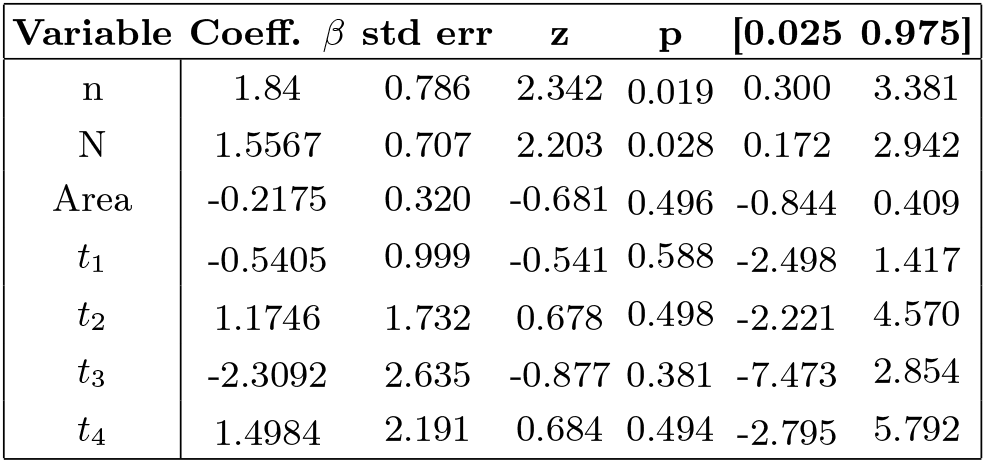
Generalized Linear Model (GLM) [40] results. (Number of samples is 96) from the statsmodel API [41]. n is the maximum recorded number of CTLs on the spheroid and N is the number of T-cells detected in the droplet, Area is the spheroid surface and *t*_1_ to *t*_4_ are the first times at which 1 to 4 CTLs are detected on the spheroid surface.

### C. Testing the statistical power of the different experimental observables on the killing

In the same chip we can record heterogeneous spheroid outcomes; some spheroids fragment very fast (complete destruction at 8 hours), whereas others are left unscathed at 14 hours (see Fig. 4e).

Since the secondary droplet contains variable CTL numbers dependent on the initial cell concentration, this leads to a range of CTL numbers in the main droplet after droplet fusion (see Fig. S1a). In addition to the number of CTLs in the droplet N, we record several other features: the first to the fourth contact times (*t*_1_ to *t*_4_, indicating the moment at which the number of cells on the spheroid goes above 1 to 4 cells), the spheroid projected area and the maximum number of CTLs on the spheroid within the 14 hours of the experiment duration n.

We conduct the test with a generalized linear model [40]. The total observation sample size is of 96 events. This test enables us to study the influence and statistical power of each variable on visible spheroid fragmentation at 14 hours. We see that only two variables have p-value below 0.05: the total number of CTLs in the well *p_n_* = 0.019, *β_n_* = 1.84) and the maximum number of CTLs on the spheroid during the time-lapse (*p_n_* = 0.028, *β_n_* = 1.56) These two measures are very correlated (see Fig. 6a); a high number of CTLs in the droplet increases the chances of having a high number of CTLs on the spheroid. The regression coefficients *β_n_* and *β_N_* were positive in both cases, confirming the positive correlation between these variables and the fragmentation probability. Interestingly, the other variables do not significantly predict the final spheroid state despite varying levels of correlation with spheroid death.

### D. Probabilistic modeling cooperative vs. independent killing of B16 cells on the spheroid by CTLs

In Fig. 5e we show that the probability of a spheroid to undergo a fragmentation depends on the maximum number of CTLs on the spheroid surface. Furthermore, the T-cell number on the spheroid is a key variable predicting the final spheroid state (see in section VI C). However, we do not know if this increase is the result of the accumulation of independent random events or if it is the signature of cooperation between T-cells. To address this question, we model the probability of a fragmentation event occurring before 14 hours with two different frameworks.

In the first model we consider that the probability for a given T-cell to cause fragmentation is independent of the presence of the other T-cells. In the second model we consider that the probability of a T-cell to produce a fragmentation event is dependent on the number of CTLs in its vicinity. In both models we use the maximum number of CTLs detected on the spheroid as a proxy for the number of interactions taking place over the duration of the experiment. This allows us to avoid including temporal aspects that would complicate the interpretation of the probabilistic models.

In quantitative terms, the probability of a fragmentation event occurring within the independent cell hypothesis is given by:

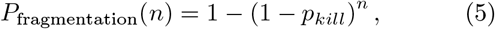

where the total number of CTLs on the spheroid is given by n and *p_kill_* is the probability to kill. Fitting equation (5) to the experimental measurements we see that the model doesn’t accurately reflect experimental results (see Fig. 4g).

A first possible improvement for the independent CTL hypothesis is proposed by Halle et al. [30] and consists in accounting for the heterogeneous nature of the CTL population. Indeed, it is well-known that CTLs differ widely in efficacy. In order to test this hypothesis we now model the CTL population as composed of independent cells, but the probability of being a fragmentation-causing CTL *k* is now itself drawn from a probability distribution 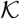 of probability denJsity *f*. The average value of *k*, 〈*k*〉 = ∫ *kf* (*k*) *dk*. We choose to not specify 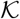 to preserve generality. Then equation 5 becomes:

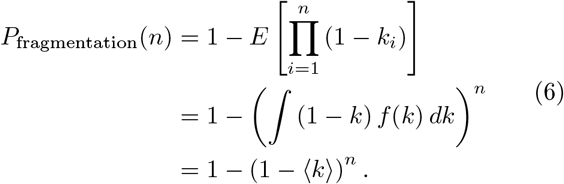

The result is similar to that in equation (5). Thus, the expected fragmentation probability is going to display a parabolic profile whatever the heterogeneity profile of the CTL population. Therefore the initial CTL population heterogeneity cannot explain the fragmentation profile recovered experimentally.

A second possible improvement to the independent CTL hypothesis is to consider that the CTLs are present on the spheroid during *T* timesteps and that they can cause a fragmentation event during any of these timesteps. Under the supplementary hypothesis that the behavior of each CTL is independent of its behavior during the previous timesteps (i.e. their behavior is *Markovian*), then the probability to see at least one fragmentation event on the spheroid during the experiment can be written as:

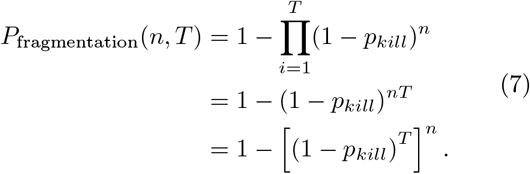

Again, the result is similar to that in equation (5) with the fragmentation probability displaying a parabolic profile. We can therefore also reject this model. In conclusion, we can reject the independent CTL hypothesis.

We will now show that a simple (though arbitrary) collective action model can recapitulate the experimental fragmentation results. In this model, we suppose that the probability for a given T-cell to lead to a fragmentation event increases when other T-cells are located nearby (i.e. on the same spheroid). The probability function *S* can be written as a sigmoid:

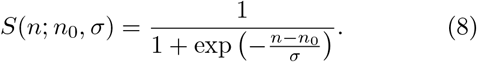

This function depends on two parameters *n*_0_ and *σ*, where *n*_0_ describes the number of neighbors required to pass a 50% probabilty and with *σ* describing how sharp is the transition. Then, the probability of witnessing fragmentation on the spheroid is given by equation 9:

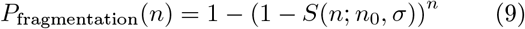

Fitting this curve to the experimental data gives a very close fit (see Fig. 4g). It is thus credible to suppose some cooperativity mechanism underlying the success of CTL killing of selected targets. However, it is not possible to deduce from these models the exact mechanism leading to target cell death.

### E. Estimating the attachment and detachment parameters (*λ_in_* and *λ_out_*)

Using the image analysis pipeline described above we are able to detect the positions of the spheroids and of the CTLs in the droplets. This information allows us to measure the number of CTLs that are attached to the spheroid at each instant. The statistics of attachment and detachment can then be used to detect whether the attraction of the cells takes place through an active process or whether it is due to random chance.

We consider a system of *N* independent CTLs undergoing Brownian-like motility in the droplet. Their motion is isotropic and the statistical properties are independent of both time and space. Their diffusion coefficient in the Matrigel is noted *D*. The spheroid itself is modeled as a sphere of diameter *a* and the droplet has a volume *V*. Each CTL can be in one of two states: in the Matrigel or on the spheroid. We consider that when a CTL encounters the spheroid it immediately attaches to it and remains attached until a detachment event occurs.

Condamin et al. [42] show that the probability of hitting the target for the first time at time *t* is:

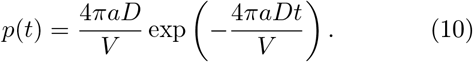

Integrating equation 10 between 0 and ∆*t* we get the probability of a cell hitting and attaching to the spheroid between times 0. and ∆*t*:

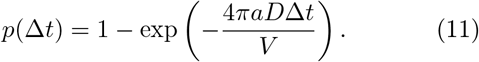

Each CTL in the Matrigel has a given probability of hitting and attaching to the spheroid during a time ∆*t*. Let us first consider the cells attaching the target over a given time-interval ∆*t*. The Matrigel droplet contains a total of *N* cells and the results in equation 11 tell us that for a single cell, the probability of attaching between times 0 and ∆*t* is:

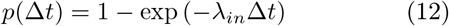

where 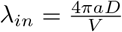 is the hitting rate. The probability of multiple independent cells hitting the target during ∆*t* follows a binomial distribution with *p* as parameter. Let us call *c* the number of contacts during ∆*t*.

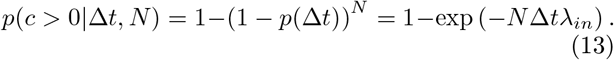

For small values of *N λ_in_*∆*t*, we see that this simplifies to:

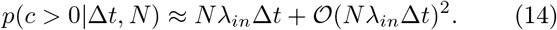

The condition *N λ_in_*∆*t* ≪ 1 means that the probability of an event happening during ∆*t* is very small, and that two or more events happening during the same time-frame is negligible.

We can reproduce the same reasoning for the probability of having one or more cells leaving the spheroid-except that the probability is now function of the number of CTLs on the spheroid *n* and not of the number of CTLs in the gel *N*.

As one can see in the model above, the attachment probability depends on a single parameter *λ_in_*. We can model the detachment dynamics of the CTLs from the spheroid in a similar manner defined by parameter *λ_out_*. *λ_in_* and *λ_out_* are the inverse of a characteristic time, and tell us how fast a cell hits the spheroid or leaves it once attached. To retrieve the parameters from the evolution of the number of CTLs attached, we count the hitting and leaving events, then we get a hitting or leaving probability *p*. Working backwards from the formula 13, we get *λ* from *p* by correcting with the number of CTLs in the ECM or on the spheroid.

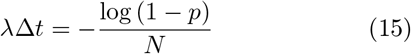

And experimentally we have 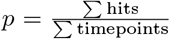 We can also specifically calculate a value of the hitting rate given a number of CTLs on the spheroid. We do so by estimating *p_n_* the probability of hitting given that there are *n* CTLs on target and using the same formula as in 15 but only counting hits and time-points where the spheroid has *n* CTLs on it.

This procedure enables us to calculate the hitting and leaving parameters of our model. To estimate the uncertainty of our estimations we conduct a bootstrapping scheme, where we randomly select experiments and calculate the hitting and leaving parameters for this subset of our initial experiment lot. The code for estimating the attachment and detachment rates from the experimental data is available upon reasonable request. From this set of parameters we retrieve a distribution for each parameter, which enables us to calculate the mean and the variance of each.

Thus, the ratio of the leaving and entering rates gives us information on the stable state distribution of cells on and off the spheroid in the droplet.

### F. Relating the spheroid fate to the number of CTLs in the droplet

We know that due to a positive feed-back loop, CTLs probability depends on a single parameter *λ_in_*. We can exhibit different accumulation rates depending on the number of CTLs on the target spheroid. We have estimated the experimental distribution of the hitting probabilities in section VI E. We also know that a certain number of CTLs on the spheroid are necessary for a high killing probability (see figure 5). Combining these two pieces of information we run the procedure in algorithm 1 which reproduces the accumulation process also described in figure 3 for a given number of CTLs in the droplet *N*.

We define the number of CTLs on the spheroid *n*_sph_, the maximum detected number of CTLs on the spheroid 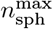 and the number of CTLs in the gel *N*_gel_. Thanks to the estimated parameters *λ_in_* and *λ_out_* we have the attachment and detachment probabilities *p*_in_ and *p*_out_. From equation 13 we get the equation for *p*_in_ (respectively *p*_out_) in equation 16:

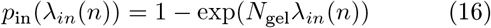

**Algorithm 1.**
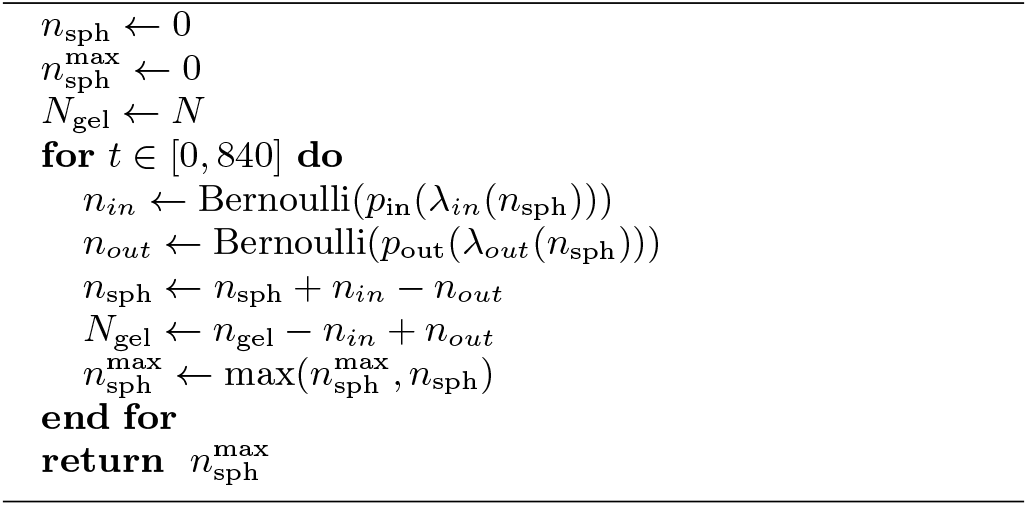
Simulate the spheroid accumulation

For a given CTL number, we simulate at each time-step the adhesion or departure of a CTL from the spheroid, with the experimentally estimated attachment or detachment probabilities. These probabilities are updated at each time step as a function of the number of CTL adhering to the spheroid. At the end of the time-loop we draw the fragmentation probability which depends on 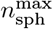 as per figure 5. By re-running this procedure 200 times for the same CTL number we get a simulated fragmentation probability that is presented in figure 6c.

